# Suspended multiwalled, acid-functionalized carbon nanotubes promote aggregation of the opportunistic pathogen *Pseudomonas aeruginosa*

**DOI:** 10.1101/2020.04.29.068973

**Authors:** Kristin Kovach, Indu Venu Sabaraya, Parth Patel, Mary Jo Kirisits, Navid B. Saleh, Vernita D. Gordon

## Abstract

The increasing prevalence of carbon nanotubes (CNTs) as components of new functional materials has the unintended consequence of causing increases in CNT concentrations in aqueous environments. Aqueous systems are reservoirs for bacteria, including human and animal pathogens, that can form biofilms. At high concentrations, CNTs have been shown to display biocidal effects; however, at low concentrations, the interaction between CNTs and bacteria is more complicated, and antimicrobial action is highly dependent upon the properties of the CNTs in suspension. Here, impact of low concentrations of multiwalled CNTs (MWCNTs) on the biofilm-forming opportunistic human pathogen *Pseudomonas aeruginosa* is studied. Using phase contrast and confocal microscopy, flow cytometry, and antibiotic tolerance assays, it is found that sub-lethal concentrations (2 mg/L) of MWCNTs promote aggregation of *P. aeruginosa* into multicellular clusters. However, the antibiotic tolerance of these “young” bacterial-CNT aggregates is similar to that of CNT-free cultures. Overall, our results indicate that the co-occurrence of MWCNTs and *P. aeruginosa* in aqueous systems, which promotes the increased number and size of bacterial aggregates, could increase the dose to which humans or animals are exposed.

## Introduction

Biofilms are communities of interacting microorganisms that are bound together in an exopolymeric matrix. Biofilm bacteria often exhibit tolerance or resistance to antibiotics and to the immune system, and the close association of bacteria within a biofilm can promote inter-cellular signaling that increases virulence.(1-5) Chronic infections caused by biofilms account for 17 million new infections and more than 0.5 million deaths in the United States each year, thereby increasing the associated healthcare costs by billions of dollars.(6) Many of these infections can be attributed to biofilm formation on medical devices, such as catheters and prosthetic heart valves.(7) As such, the formation of anti-biofilm surfaces is of great interest to scientists and healthcare providers. Carbon nanotubes (CNTs) incorporated into polymer composites have been shown to kill bacteria and therefore reduce biofilm growth for several strains of bacteria; in particular, a concentration of ∼3-5% by weight of CNTs in a solid composite material is typically lethal to 80-90% of the bacteria present.(8, 9) However, when CNTs are at low concentrations in aqueous suspension, the interaction between CNTs and bacteria becomes more complex. Some have suggested that low concentrations of suspended SWCNTs might provide a net benefit to bacterial populations.(8, 10-13)

In addition to their antimicrobial activity, CNTs can enhance the strength and conductivity of composites and, therefore, are used increasingly to develop new materials.(10, 14) As the manufacture and application of CNT-containing composites increase, the release of CNTs to natural and engineered water systems also is likely to increase; this could be the result of CNT discharge as by-products of the manufacturing process or CNT release as functionalized materials that are utilized, discarded, and then environmentally degraded.(15) Models have been used to estimate concentrations of nanomaterials, including CNTs, in the environment.(15-18) For instance, models approximated that there were 0.001 ng/L of CNTs in surface water in the United States in 2008. CNT concentrations were expected to increase annually by 46 ng/kg in sediment and 0.56 ng/kg in soil; comparable values are estimated for Europe, and all of these values are predicted to grow as the usage of CNTs becomes more widespread.(18)

Although some studies have examined the effects of low-concentration suspensions of SWCNTs on microbial inactivation, little is known about how sub-lethal concentrations of suspended MWCNTs impact bacteria.(11, 13, 19, 20) *Pseudomonas aeruginosa* is a highly-studied, opportunistic human pathogen that readily forms biofilms that can cause deadly chronic infections.(21) *P. aeruginosa* also is widespread in natural and engineered environments, including waterways and water treatment systems.(22)

Here, we show that sub-lethal concentrations of suspended MWCNTs promote the formation of large *P. aeruginosa* aggregates that contain MWCNTs. Previous work by others has shown that spontaneously-formed, CNT-free, suspended aggregates of *P. aeruginosa* have many characteristics of biofilms, including increased antibiotic resistance and virulence.(23) Our own earlier work has demonstrated that bacterial aggregates can be more effective in initiating biofilm growth on surfaces than are single cells.(24) This raises the question as to whether increasing the environmental load of suspended CNTs could increase the likelihood that humans and other animals would be exposed to an infectious dose of bacteria in a concentrated, biofilm-like, antibiotic-resistant state. However, in partial amelioration of this concern, we find that MWCNT-containing bacterial aggregates are no more tolerant of antibiotics than are bacterial aggregates that do not contain CNTs.

## Materials and Methods

### Bacterial Cultures

The bacterial strain used in this study was *P. aeruginosa* PAO1 that constitutively expresses the green fluorescent protein (GFP).(25) GFP was used in flow cytometry and microscopy analyses.

### Acquisition and Functionalization of CNTs

MWCNTs with outer diameters of 8-15 nm were purchased from CheapTubes (Cambridgeport, VT) and functionalized using a previously-published procedure.(26) In brief, to etch oxygenated functional groups (e.g., -OH, -COOH, -COH)(27, 28) on the MWCNTs, 1 g of nanotubes was added to a mixture of concentrated nitric and sulfuric acid in a round-bottomed flask. The mixture was sonicated to disperse MWCNTs and refluxed at 100 °C for 3 h under continuous stirring. The oxidized MWCNTs were subsequently filtered until the pH of the filtrate reached >5.5 and then were dried for 48 h in a desiccator.

### Bacterial Media

Davis Minimal Medium (DMM) was used for growing bacteria in liquid suspension. It consists of a solution in Millipore water containing 1 g/L ammonium sulfate, 7 g/L dipotassium phosphate, 2 g/L monopotassium phosphate, 0.5 g/L sodium citrate, and 0.1 g/L magnesium sulfate, supplemented with a filter-sterilized solution of dextrose to a final concentration of 1 g/L (where all materials except water were purchased from Sigma-Aldrich). DMM is autoclaved and allowed to cool before use.

Bacterial plate counts were performed on lysogeny broth (LB-Miller) agar plates (10 g tryptone, 5 g yeast extract, 10 g NaCl, and 15 g agar (all from Sigma-Aldrich) in 1 L of Millipore water). Ten-microliter aliquots of diluted bacterial samples were spot-plated in duplicate on the agar plates and incubated for 24 h at 37 °C before counting.

### Bacterial Culturing

*P. aeruginosa* PAO1 was cultured overnight (∼10 h) in DMM at 37 °C with shaking to reach the exponential phase. The cells were subcultured, such that 40 µL of the overnight culture was added to 4 mL of fresh DMM, yielding a final concentration of about 2 × 10^7^ colony-forming units (CFU)/mL. One set of subcultures was supplemented with MWCNTs (400 µL of 20 mg/L MWCNTs suspended in DMM was added to each subculture), and another set of subcultures was maintained as controls (400 µL of sterile DMM was added to each subculture). These cultures were used for measurements of growth, dead staining, and antibiotic tolerance and for microscopy of aggregate structure as described in the following sections.

### Growth Measurements

Every 1-2 h, one control subculture (containing no MWCNTs) and one MWCNT- containing subculture were sacrificed. Here, cultures were sonicated for 5-10 seconds to break up aggregates, and aliquots were removed for measurement of optical density at 600 nm (OD_600_) and for plate counts. Aliquots (100-µL) were used for OD_600_ measurements with a Genesys spectrophotometer (Thermo Fisher Scientific, Waltham, MA). Plate counts were performed as described in an earlier section. Three biological replicates, grown and interrogated on different days, were used for OD_600_ measurements; two biological replicates, also grown and interrogated on different days, were used for plate counts.

### Dead Staining and Image Analysis of Bacterial Aggregates

After the subcultures had been incubated for 5 h in the absence or presence of MWCNTs, the OD_600nm_ of each sample was measured, and 3 µL of the DNA-stain propidium iodide (PI) was added to 1 mL of culture. When a bacterial cell dies and its membrane is compromised, the intracellular DNA binds to PI to identify a dead cell.

A 20-μL aliquot of each PI-stained sample was imaged via confocal laser scanning microscopy (CLSM) using an Olympus FV1000 motorized inverted IX81 microscope (Olympus, Center Valley, PA) with a 100× oil-immersion objective; 488 nm and 543 nm lasers were used for GFP and PI excitation, respectively. Images were captured using FV10-ASW version 3.1 software (Olympus, Center Valley, PA). Three to four biological replicates were evaluated for the control and MWCNT-containing cultures; for each replicate, three to four regions were selected at random, and confocal z-stacks of each region were acquired.

To analyze confocal micrographs of PI-stained samples, ImageJ was first used to create a maximum-intensity projection from each z-stack, and then a threshold brightness value was applied to the resulting two-dimensional images. This produced a binary (bright or dark) image, and the particle analysis feature was used to measure the sizes of bright areas (identified by ImageJ as “particles” irrespective of size, so an aggregate containing 100 bacteria would count as one particle) in both the GFP and PI channels. Next, in Matlab, these sizes were used to sort images into those containing both single bacteria and aggregates and those containing only aggregates. For each image, a Matlab code was used to count the number of particles that were bright with GFP fluorescence and the number of particles that were bright with PI fluorescence; this count included both single cells and aggregates, when present. The ratio of the number of GFP-bright particles to the number of PI-bright particles was measured for each image and comparison of this ratio among different images was used to estimate differences in the proportion of live and dead bacteria.

### Antibiotic Tolerance Testing with Confocal Laser Scanning Microscopy

To test for antibiotic tolerance in the subcultures, after four hours of subculture growth in the presence or absence of MWCNTs, tobramycin was added to each subculture to a final concentration of 10 or 20 μg/mL and incubated for one hour. Then, OD_600nm_ was measured and 3 μL of PI was added to 1 mL of each tobramycin-treated subculture. Imaging and analysis were conducted as described in the previous section.

### Microscopy Imaging of Aggregates

To assess aggregates at different time points during subculture growth, an aliquot of subculture was removed every 1-2 h and was imaged using phase contrast microscopy and epifluorescence microscopy on an Olympus IX71 inverted microscope with a 60× objective and GFP filter cube (Olympus, Center Valley, PA), QImaging Exi Blue CCD camera, and QCapture Pro 6 software (Teledyne Qimaging, Surrey, British Columbia, Canada).

### Flow Cytometry of Bacterial Aggregates

Flow cytometry was used for high-throughput measurements of aggregate development. To 4 mL fresh DMM, we added one of the following: 40 μL from an overnight *P. aeruginosa* culture plus 400 μL of sterile DMM; 40 μL from an overnight culture of *P. aeruginosa* plus 400 μL of MWCNTs suspended in DMM; or 400 μL of DMM. The resulting samples were incubated for 3 h with shaking at 37 °C. Then, cultures were vortexed for about 5 seconds to resuspend the material that had settled out, and the OD_600nm_ of the cultures was measured. Samples were placed into Eppendorf tubes for flow cytometry on a BD Accuri Flow Cytometer (BD Biosciences, San Jose, CA), mixing the contents by inverting the tubes 10× before each run. For each measurement, the flow cytometer was run for 2.5 min at a flow rate of 40 μL/min with a threshold for events larger than 80,000 forward scatter height. Two independent biological replicates were tested under each experimental condition; each biological replicate had three technical replicates for bacteria cultured without MWCNT and two technical replicates for bacteria cultured with MWCNT.

### Fitting and statistical analysis

Data fitting and statistical analyses were performed in Excel (Microsoft, Redmond, WA, USA). Specifically, we used the built-in one-way analysis of variance (ANOVA) test with a p-value of 0.05 used as the threshold for significance.

## Results and Discussion

OD_600nm_ and CFU counts were used to assess how suspended MWCNTs impacted bacterial growth (Fig 1A). We fit exponential functions (of the form *N*(*t*) = *N*(0)*e*^*kt*^) to the exponential-growth region of each growth curve, where *t* is time and *k* is the first-order rate constant (Fig 1B). Eight out of ten of these fits had an R^2^ value greater than 0.9, indicating that growth is well-described by this expression. It was not unexpected that the rate constants calculated with OD_600nm_ and CFU/mL data were different from one another because OD_600nm_ measures the scattering of light, which is affected by particle shape and size, among other factors. Single-factor ANOVA analysis of both the OD_600nm_ and CFU/mL data indicates no statistically significant difference in growth between subcultures with and without suspended MWCNTs; p-values are 0.76 for OD measurements and 0.9 for CFU measurements. Therefore, we conclude that the type and concentration of MWCNTs used in these experiments do not impact bacterial growth rate.

**Fig 1.**
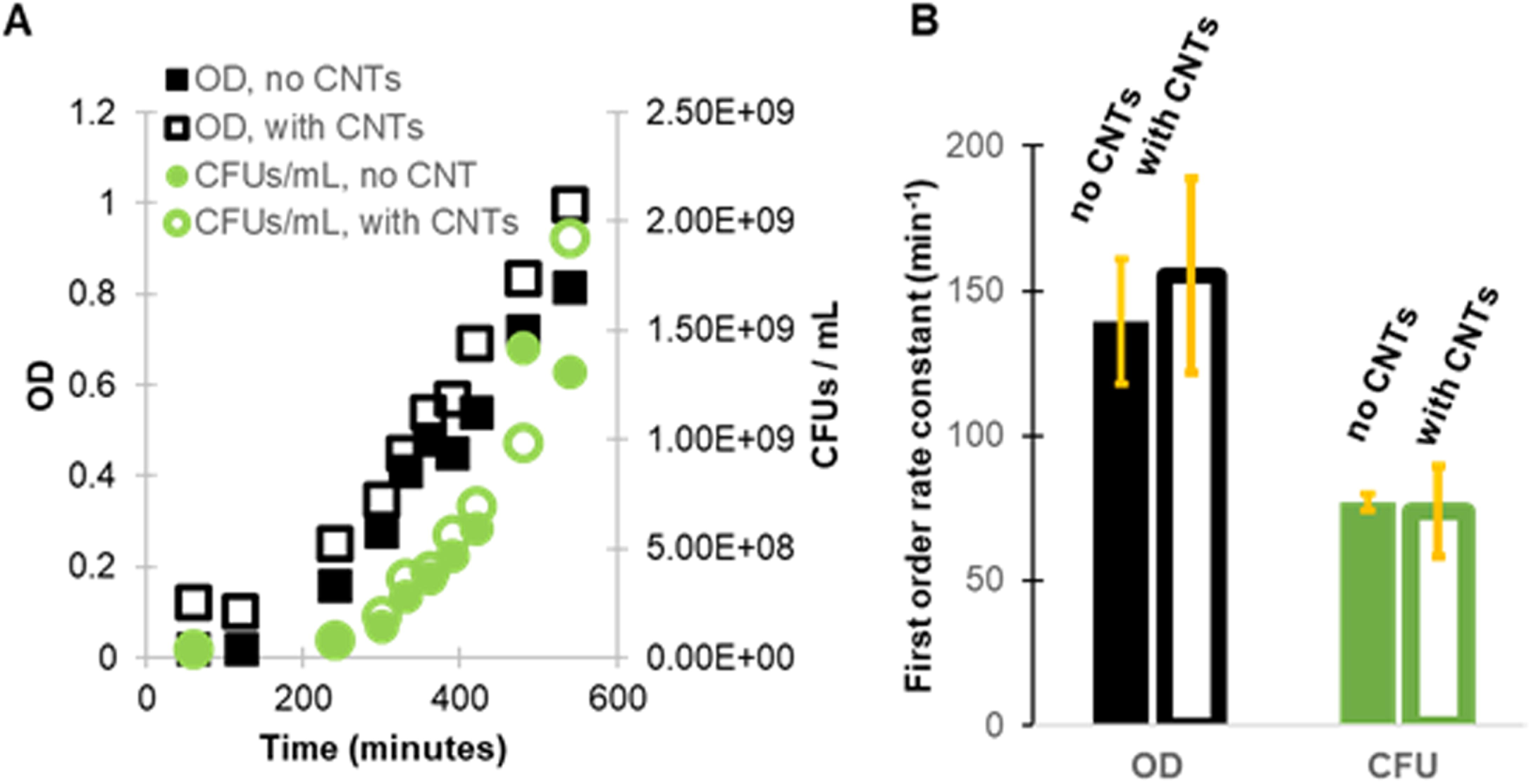
At the concentration used in this study, CNTs do not significantly impact the growth rate of *P. aeruginosa*. (A) Representative growth measurements over 9 h, using OD_600_ and CFU/mL, with and without MWCNTs. (B) First-order rate constants calculated by fitting exponential functions to growth data such as those shown in (A). Three biological replicates (N=3), grown and interrogated on different days, were used for OD_600_ measurements, and two biological replicates (N=2), also grown and interrogated on different days, were used for plate counts. Error bars are the standard error of the mean.

However, to the naked eye, bacterial subcultures grown in the presence of MWCNTs look strikingly different from subcultures grown in the absence of MWCNTs. Namely, the MWCNT-containing subcultures contain dark clumps that are not observed in either the MWCNT-free subcultures or in bacteria-free suspensions of MWCNTs (Supplementary Fig S1). *P. aeruginosa* grown in liquid culture is known to spontaneously form aggregates even in the absence of MWCNTs(23, 24), but such aggregates are not visible to the naked eye (Supplementary Figure S1A). Phase-contrast micrographs of bacteria-CNT aggregates show that the bacteria surround a dark core, presumably of clumped MWCNTs, but dark cores are not observed in the aggregates that form in CNT-free cultures (Fig 2).

**Fig 2.**
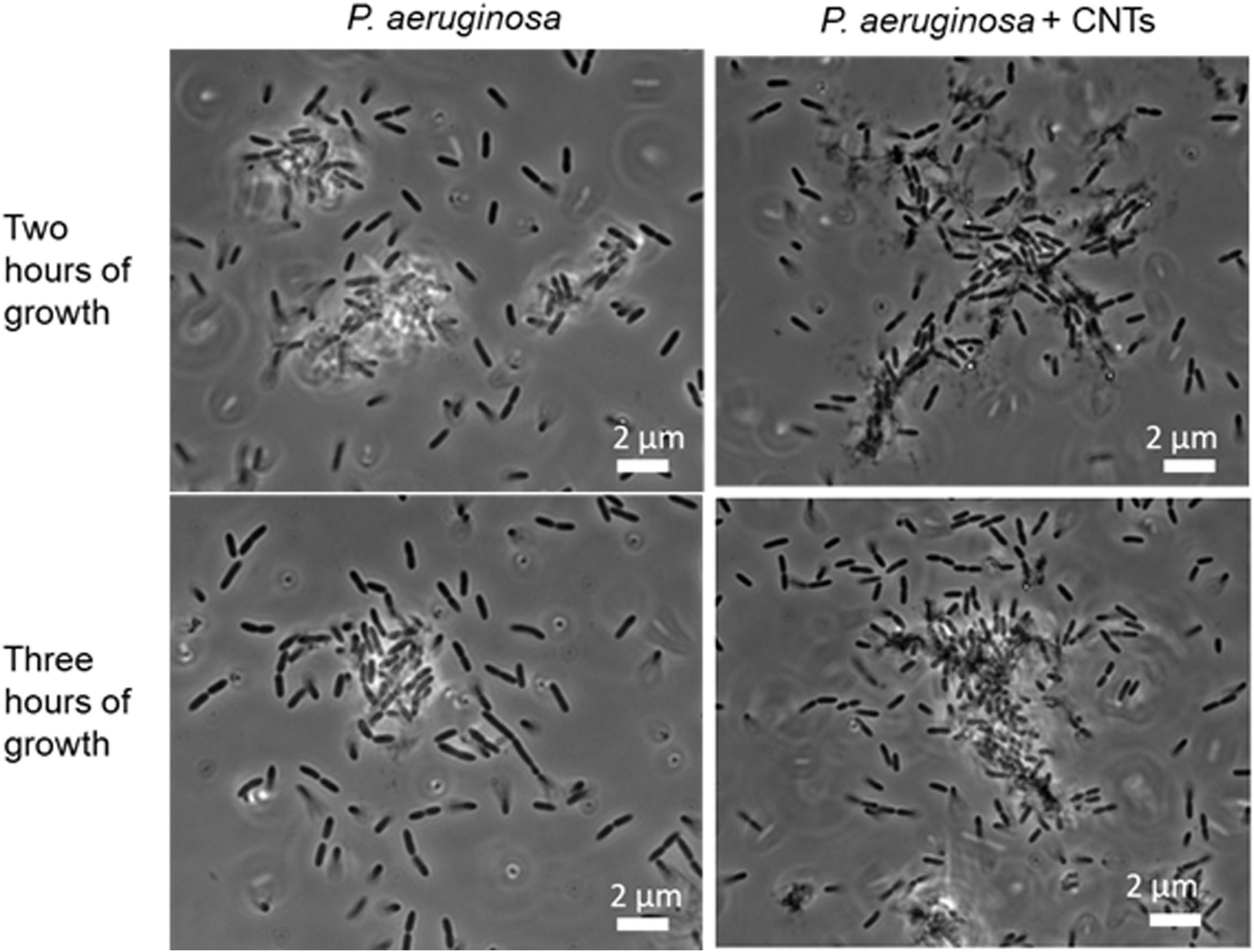
Phase-contrast micrographs of *P. aeruginosa* in the presence or absence of CNTs. Aggregates that form in the presence of MWCNTs (right) have a denser and more-elongated structure than do bacterial aggregates that form in the absence of MWCNTs (left). Aggregates formed in the presence of MWCNTs have dark cores that we attribute to clustered MWCNTs.

To quantify the impact of MWCNTs on the number and size of *P. aeruginosa* aggregates, we use flow cytometry. We define a gate in terms of both high scatter area (SSC-A) and high GFP fluorescence area (FL1-A), which corresponds to objects that are much larger than single bacteria in two perpendicular directions and that are made up of GFP-expressing bacteria. We use the same gate for all measurements, and we find that the fraction of the bacterial sample within that gate is consistently much larger in the subcultures containing MWCNTs than in the subcultures that do not contain MWCNTs (Fig 3). These results indicate that there are approximately 2.5× more aggregates present in the samples with MWCNTs, which is consistent with the idea that MWCNTs promote aggregate formation. Single-factor ANOVA analysis yields a p-value of 10^−5^ when all sample types (including those without bacteria) are included and a p-value of 0.008 when only samples containing bacteria grown with and without MWCNTs are compared; these results indicate statistical significance.

**Fig 3.**
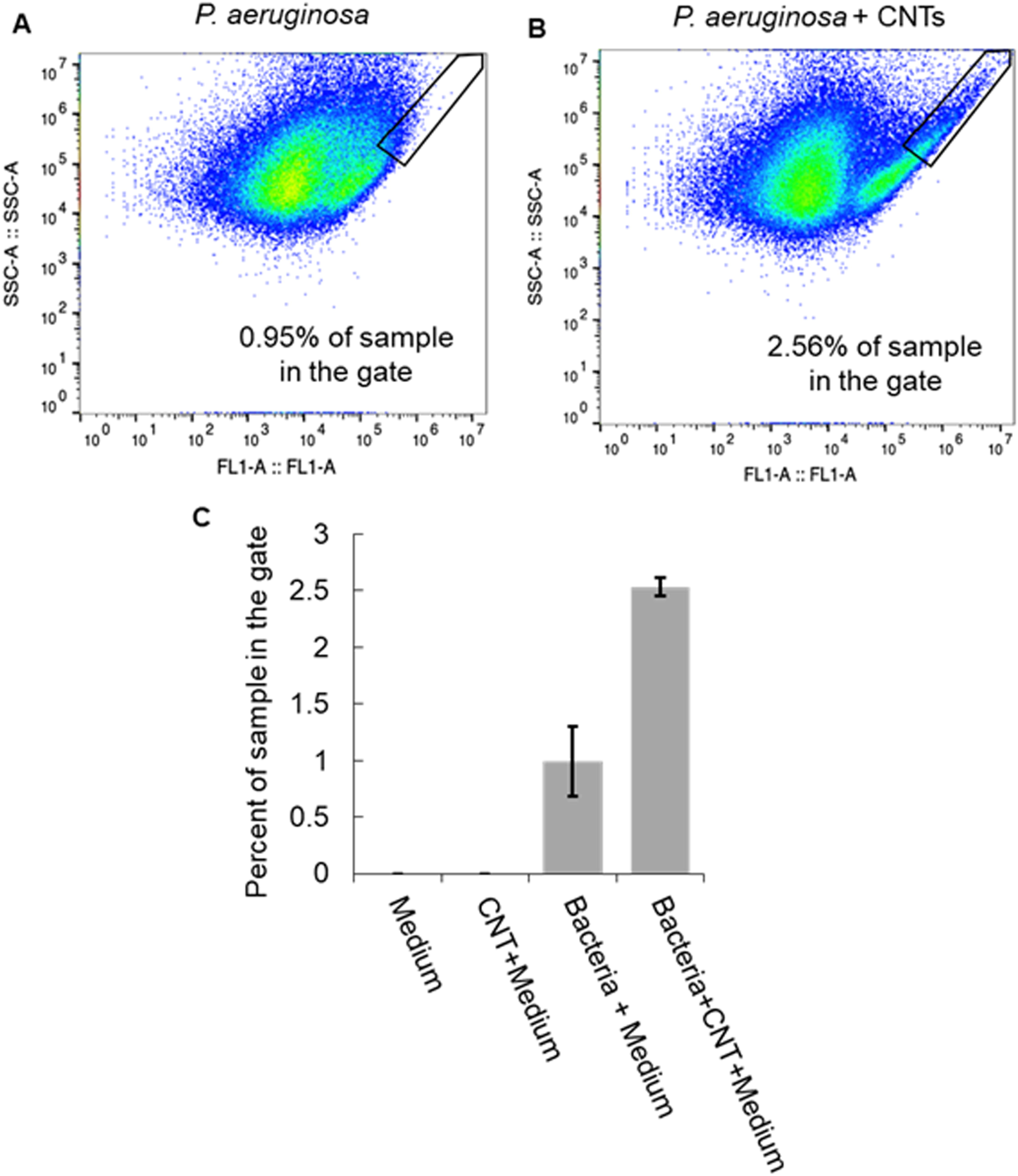
Flow cytometry is used to measure the percentage of each subculture containing large aggregates. Examples of data from flow cytometry containing (A) *P. aeruginosa* only and (B) *P. aeruginosa* with MWCNTs. The vertical axes, labeled SSC-A, give the side-scatter area, and the horizontal axes, labeled FL1-A, give the GFP fluorescence area. Black lines define a gate with both high side-scatter and high fluorescence areas. This region is the gate we use to measure the number of very large fluorescent objects in the sample. (C) For three separate sets of samples and flow cytometry trials (N=3), we find that bacterial subcultures containing MWCNTs have, on average, more than twice as many large aggregates than do MWCNT-free bacterial subcultures. No large aggregates were found in the sterile medium or the sterile medium containing MWCNTs. Error bars are the standard error of the mean.

We have previously shown that pre-formed aggregates of bacteria can accelerate and dominate the growth of biofilms on the surface to which aggregates attach.(24) Others have reported that bacterial aggregates themselves can have biofilm-like properties, such as the activation of quorum sensing, antibiotic resistance, and resistance to phagocytic clearance by the immune system.(23) Therefore, the prospect of inadvertently promoting the formation of more and larger bacterial aggregates through the presence of MWCNTs in the natural environment or in water treatment systems is concerning. Of these, antibiotic resistance might be the biggest concern from a public health standpoint. To evaluate the degree to which bacteria tolerate antibiotics, we treat cultures with tobramycin, which is an aminoglycoside antibiotic that is the front-line drug for treating some *P. aeruginosa* infections.(29) Using dead staining and quantitative analysis of confocal microscopy images, we compare MWCNT-containing and MWCNT-free subcultures to assay the degree to which greater aggregation, promoted by MWCNTs, is associated with greater antibiotic tolerance (Fig 4A,B). Contrary to our expectations, we find that there is no statistically significant difference in antibiotic tolerance between MWCNT-containing and MWCNT-free cultures, regardless of whether a mixture of single cells and aggregates or only aggregates are being analyzed (Fig 4C,D). Given that the bacterial-CNT aggregates were developed only over 4 h, these might be considered “young” biofilms. If the bacterial-CNT aggregates were allowed to develop over longer periods of time (i.e., becoming “mature” biofilms), these might show increased antibiotic tolerance as compared to bacteria-only cultures. This is supported by the work of Høiby et al., who demonstrated that the minimum biofilm eradication concentration of an antibiotic increased as biofilm age increased (i.e., planktonic cells to young biofilm to mature biofilm).(30) Additionally, suspended bacterial aggregates formed without CNTs were found to have greater antibiotic tolerance than planktonic bacteria.(23) However, there is some precedent for similar tolerances to nanomaterial insults between mature biofilm and planktonic populations; for *Escherichia coli* and *P. aeruginosa*, biofilm and planktonic cells were shown to have similar tolerance to silver nanoparticles.(31) Thus, fundamental studies of stress in bacteria (both biofilm and planktonic cells) due to nanomaterial exposures are needed to better understand the impact of nanomaterials on bacteria at the cellular level.

**Fig 4.**
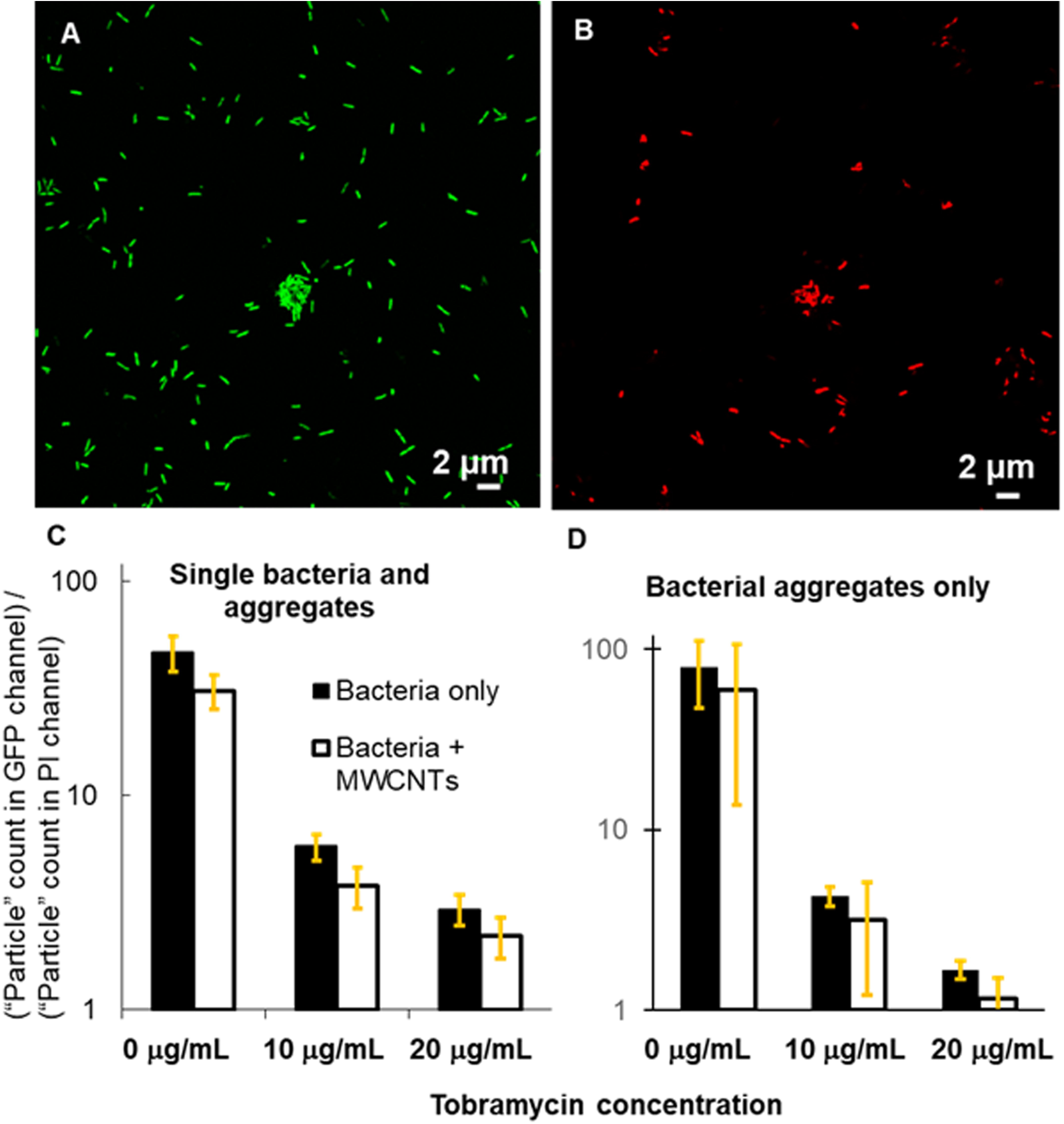
Results from dead-staining of *P. aeruginosa* cultures. Microscopy images show single bacteria and aggregates after exposure to tobramycin at 20 μg/mL in (A) the GFP fluorescence channel and (B) the PI fluorescence channel (dead). (C and D) For the three tobramycin concentrations tested, for both (C) single bacteria and aggregates and (D) aggregates only, the surviving fraction of the population is slightly less for the MWCNT- containing sample. However, single-factor ANOVA analysis shows that the differences between the MWCNT-containing samples and the corresponding MWCNT-free samples are not statistically significant. For aggregates and single cells imaged together, five replicate experiments were done for MWCNT-free samples and four for MWCNT- containing samples; for aggregates imaged without single cells, three replicate experiments each were conducted. Error bars are the standard error of the mean.

## Conclusions

Here, we show that the opportunistic human pathogen *P. aeruginosa*, when in the presence of a sub-lethal concentration of suspended MWCNTs, uses the MWCNTs as a scaffold to build more large multicellular aggregates than it could in the absence of MWCNTs. As MWCNTs unintentionally become more widespread in natural and engineered water systems, the occurrence of such bacterial-CNT aggregates presents the possibility that humans and other animals might be exposed to a high-density dose of bacteria.

However, we find that our “young” bacterial-CNT aggregates (developed over four hours in MWCNT-containing bacterial cultures) do not have greater antibiotic tolerance than do CNT-free cultures. Our data suggest that, if caught early, infections that begin as “young” MWCNT-based aggregates might be amenable to standard antibiotic treatment. Additional work is needed to determine whether the antibiotic tolerance of mature bacterial-CNT aggregates exceeds that of single planktonic cells.

Others have shown that SWCNTs and MWCNTs, in sufficient concentration, are promising additives to composite materials for creating antimicrobial surfaces.(8, 9, 12, 13) However, the manufacture and use of such composites likely will serve to increase CNT concentrations in natural and engineered water systems. The work we present here suggests that the advantages of MWCNT-functionalized materials must be weighed against the potential disadvantage of inadvertently increasing the number and size of bacterial aggregates, which could increase the dose to which humans and animals might be exposed.

## Acknowledgements

This work was funded by grants from the National Science Foundation (CBET 0846719 to MJK; CBET 1916709 to MJK and NBS; CMMI 1727544 to VDG), the Cystic Fibrosis Foundation (GORDON1710 to VDG), and the National Institutes of Health (5 R01 AI121500 to VDG). KK was the recipient of a Graduate Research Fellowship from the National Science Foundation for part of the time this work was done (DGE-1110007). GFP-expressing PAO1 was kindly provided by Prof. Matthew Parsek, University of Washington, Seattle. Flow cytometry measurements were done at the Center for Biomedical Research Support, College of Natural Sciences, University of Texas at Austin.

## Supporting Information Caption

**Supplementary Figure S1**. (A) When bacteria are grown in the presence of suspended MWCNTs, dark clumps appear in the culture after several hours of growth (right, *P. aeruginosa* + MWCNTs). Such clumps are not observed in a MWCNT-free culture grown under the same conditions (left, *P. aeruginosa*). (B) Because MWCNTs are functionalized with oxygenated groups (-OH, -COOH, -COH) to prevent aggregation, suspensions of MWCNTs without bacteria do not contain visible clumps even at high concentrations of MWCNTs.

## References

1. Bjarnsholt T, Jensen PO, Fiandaca MJ, Pedersen J, Hansen CR, Anderson CB, et al. Pseudomonas aeruginosa biofilms in the respiratory tract of cystic fibrosis patients. Pediatric Pulmonology. 2009;44(6):547–58.

2. Fexby S, Bjarnsholt T, Jensen PØ, Roos V, Høiby N, Givskov M, et al. Biological Trojan Horse: Antigen 43 Provides Specific Bacterial Uptake and Survival in Human Neutrophils. Infection and Immunity. 2007;75(1):30–4.

3. Bjarnsholt T, Jensen PØ, Burmølle M, Hentzer M, Haagensen JAJ, Hougen HP, et al. Pseudomonas aeruginosa tolerance to tobramycin, hydrogen peroxide and polymorphonuclear leukocytes is quorum-sensing dependent. Microbiology. 2005;151(2):373–83.

4. Wessel AK, Arshad TA, Fitzpatrick M, Connell JL, Bonnecaze RT, Shear JB, et al. Oxygen Limitation within a Bacterial Aggregate. mBio. 2014;5(2):e00992–14.

5. Walters MC, Roe F, Bugnicourt A, Franklin MJ, Stewart PS. Contributions of Antibiotic Penetration, Oxygen Limitation, and Low Metabolic Activity to Tolerance of <em>Pseudomonas aeruginosa</em> Biofilms to Ciprofloxacin and Tobramycin. Antimicrobial Agents and Chemotherapy. 2003;47(1):317–23.

6. Romling U, Kjelleberg S, Normark S, Nyman L, Uhlin B, Akerlund B. Microbial Biofilm Formation: a need to act. J Intern Med. 2014;276(2):98–110.

7. Chen M, Yu Q, Sun H. Novel Strategies for the Prevention and Treatment of Biofilm Related Infections. Int J Mol Sci. 2013;14(9):18488–501.

8. Goodwin DG, Marsh K, Sosa I, Payne J, Gorham J, Bouwer E, et al. Interactions of Microorganisms with Polymer Nanocomposite Surfaces Containing Oxidized Carbon Nanotubes. Environmental Science & Technology. 2015;49(9):5484–92.

9. Ahmed F, Santos C, Vergara R, Tria M, Advincula R, Rodrigues D. Antimicrobial Applications of Electroactive PVK-SWNT Nanocomposites. Environ Sci Technol. 2012;46(3):1804–10.

10. Goodwin DG, Xia Z, Gordon T, Gao C, Bouwer E, Fairbrother D. Biofilm development on carbon nanotube/polymer nanocomposites. Environmental Science: Nano. 2016(3):545–58.

11. Rodrigues DF, Elimelech M. Toxic Effects of Single-Walled Carbon Nanotubes in the Development of <i>E. coli</i> Biofilm. Environmental Science & Technology. 2010;44(12):4583–9.

12. Kang S, Mauter MS, Elimelech M. Physiochemical determinants of Multiwalled Carbon Nanotube Bacterial Cytotoxicity. Environ Sci Technol. 2008;42(19):7528–34.

13. Yang C, Mamouni J, Tang Y, Yang L. Antimicrobial activity of single-walled carbon nanotubes: Length effect. Langmuir. 2010;26(20):16013–9.

14. Moniruzzaman M, Winey KI. Polymer Nanocomposites Containing Carbon Nanotubes. Macromolecules. 2006;39(16):5194–205.

15. Nowack B, Bucheli TD. Occurrence, behavior and effects of nanoparticles in the environment. Environmental Pollution. 2007;150(1):5–22.

16. Petersen EJ, Henry TB. Methodological Considerations for Testing the Ecotoxicity of Carbon Nanotubes and Fullerenes: Review. Environmental Toxicology and Chemistry. 2012;31(1).

17. Petersen EJ, Zhang L, Mattison NT, O’Carroll DM, Whelton AJ, Uddin N, et al. Potential Release Pathways, Environmental Fate, And Ecological Risks of Carbon Nanotubes. Environ Sci Technol. 2011;45(23):9837–56.

18. Gottschalk F, Sonderer T, Scholz RW, Nowack B. Modeled Environmental Concentrations of Engineered Nanomaterials (TiO2, ZnO, Ag, CNT, Fullerenes) for Different Regions. Environ Sci Technol. 2009;43(24):9216–22.

19. Arias LR, Yang L. Inactivation of bacterial pathogens by carbon nanotubes in suspensions. Langmuir. 2009;25(5):3003–12.

20. Mohanty A, Wei L, Lu L, Chen Y, Cao B. Impact of Sublethal Levels of Single-Wall Carbon Nanotubes on Pyoverdine Production in Pseudomonas aeruginosa and Its Environmental Applications. Environ Sci Technol Lett. 2015;2(4):105–11.

21. Alhede M, Bjarnsholt T, Givskov M, Alhede M. Pseudomonas aeruginosa Biofilms: Mechanisms of Immune Evasion. 1 ed: Copyright © 2014 Elsevier Inc. All rights reserved.; 2014. 1–40 p.

22. Green SK, Schroth MN, Cho JJ, Kominos SD, Vitanza-Jack VB. Agricultural Plants and Soil as a Reservoir for Pseudomonas aeruginosa. Appl Microbiol. 1974;28(6):987–91.

23. Alhede M, Kragh KN, Qvortrup K, Allesen-Holm M, van Gennip M, Christensen LD, et al. Phenotypes of Non-Attached Pseudomonas aeruginosa Aggregates Resemble Surface Attached Biofilm. PLoS ONE. 2011;6(11):e27943–e.

24. Kragh KN, Hutchison JB, Melaugh G, Rodesney C, Roberts AE, Irie Y, et al. Role of Multicellular Aggregates in Biofilm Formation. mBio. 2016;7(2):e00237–16.

25. B W Holloway a, Morgan AF. Genome Organization in Pseudomonas. Annual Review of Microbiology. 1986;40(1):79–105.

26. Das D, Plazas-Tuttle J, Sabaraya IV, Jain SS, Sabo-Attwood T, Saleh NB. An elegant method for large scale synthesis of metal oxide–carbon nanotube nanohybrids for nano-environmental application and implication studies. Environmental Science: Nano. 2017;4(1):60–8.

27. Avilés F, Cauich-Rodríguez JV, Moo-Tah L, May-Pat A, Vargas-Coronado R. Evaluation of mild acid oxidation treatments for MWCNT functionalization. Carbon. 2009;47(13):2970–5.

28. Gong H, Kim S-T, Lee JD, Yim S. Simple quantification of surface carboxylic acids on chemically oxidized multi-walled carbon nanotubes. Applied Surface Science. 2013;266:219–24.

29. Takahashi Y, Igarashi M. Destination of aminoglycoside antibiotics in the ‘post-antibiotic era’. J Antibiot. 2018;71(1):4–14.

30. Høiby N, Henneberg K-Å, Wang H, Stavnsbjerg C, Bjarnsholt T, Ciofu O, et al. Formation of Pseudomonas aeruginosa inhibition zone during tobramycin disk diffusion is due to transition from planktonic to biofilm mode of growth. International Journal of Antimicrobial Agents. 2019;53(5):564–73.

31. Saleh NB, Chambers B, Aich N, Plazas-Tuttle J, Phung-Ngoc HN, Kirisits MJ. Mechanistic lessons learned from studies of planktonic bacteria with metallic nanomaterials: implications for interactions between nanomaterials and biofilm bacteria. Frontiers in Microbiology. 2015;6(677).

