## Supplementary figures and images for "Suspended multiwalled, acid-functionalized carbon nanotubes promote aggregation of the opportunistic pathogen *Pseudomonas aeruginosa*"

### Supplementary Figure S1

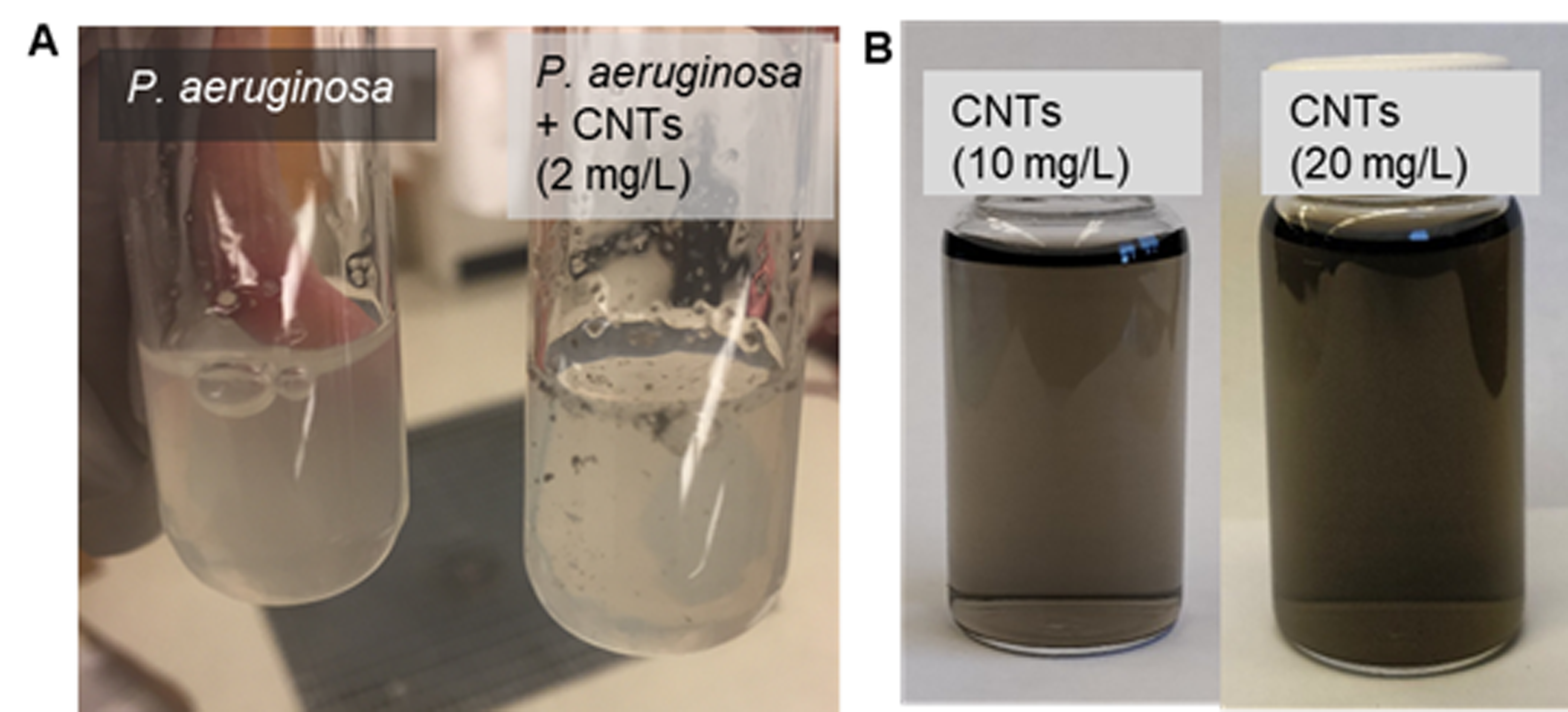
